# A diagnostic support system based on interpretable machine learning and oscillometry for accurate diagnosis of respiratory dysfunction in silicosis

**DOI:** 10.1101/2025.01.08.632001

**Authors:** Jorge Luís Machado do Amaral, Cíntia Moraes de Sá Sousa, Caroline de Oliveira Ribeiro, Paula Morisco de Sá, Agnaldo José Lopes, Pedro Lopes de Melo

## Abstract

Silicosis, the most dangerous and common lung illness associated with breathing in mineral dust, is a significant health concern. Spirometry, the traditional method for evaluating pulmonary functions, requires high patient compliance. Respiratory Oscillometry and electrical models are being studied to evaluate the respiratory system. This study aims to harness the power of machine learning (ML) to enhance the accuracy and interpretability of oscillometric parameters in silicosis. The data was obtained from 109 volunteers (60 in the training and 49 in the validation groups). Some supervised ML algorithms were selected for tests: K-Nearest Neighbors, Logistic Regression, Random Forest, CatBoost (CAT), Explainable Boosting Machines (EBM), and a deep learning algorithm. Two synthetic data generation algorithms were also applied. Initially, this study revealed the most accurate oscillometric parameter: the resonant frequency (fr, AUC=0.86), indicating a moderate accuracy (0.70-0.90). Next, original oscillometric parameters were used as input in the selected algorithms. EBM (AUC=0.93) and HyperTab (AUC=0.95) demonstrated the best performance. When feature selection was applied, HyperTab (AUC=0.94), EBM (AUC=0.94), and Catboost (AUC=0.93) emerged as the most accurate results. Finally, external validation resulted in a high diagnostic accuracy (AUC=0.96). Machine learning algorithms introduced enhanced accuracy in diagnosing respiratory changes associated with silicosis. The HyperTab and EBM achieved a high diagnostic accuracy range, and EBM explains the importance of the features and their interactions. This AI-assisted workflow has the potential to serve as a valuable decision support tool for clinicians, which can enhance their decision-making process, ultimately leading to improved accuracy and efficiency.

## INTRODUCTION

The oldest, most dangerous, and most common lung illness associated with breathing in mineral dust is silicosis. This disorder is one of the most common occupational diseases globally and can arise in various jobs, including cement manufacturing, ceramic and metal industries, shipbuilding, and maintenance [1, 2]. According to Leung, Yu, and Chen [1], radiological findings are typically used to identify this illness, while Cox, Rose, and Lynch [3] have suggested that high-resolution computed tomography is the most effective technique for evaluating respiratory disorders caused by the environment and at work. Spirometry is a frequent tool used to evaluate pulmonary function in silicosis patients. However, this procedure necessitates high patient compliance, as patients must execute maximal forced expiratory maneuvers, which may alter airway features due to changes in bronchial tone. Furthermore, some patients cannot perform these exams due to the need for the cited difficult maneuvers [4].

System Identification aims to build detailed mathematical models of dynamic systems based on their input-output relationships. One approach that has been evaluated to examine the respiratory system biomechanics is the forced oscillation technique (FOT), also known as respiratory oscillometry [5, 6]. It is a specific case of system identification, in which the characteristics of a system (in this case, the respiratory system) are determined based on how the system responds to an external input (pressure waves) [7]. The method is noninvasive and assesses the respiratory system resistance and reactance using exams performed under spontaneous ventilation. These are practical advantages in occupational settings where individuals may hesitate to undergo more invasive and complex tests.

Linking detailed oscillometry data with electrical models offers a powerful way to understand respiratory diseases better, enabling clinicians and researchers to analyze the intricate relationships between airflow, pressure, and lung mechanics with greater detail [7]. The extended Resistance-Inertance-Compliance (eRIC) model enhances sensitivity in detecting lung function abnormalities, facilitating the distinction between obstructed and non-obstructed airways [8, 9], enabling early identification of pulmonary diseases [10, 11], and providing a quantitative assessment of lung function impairment [12].

As observed in the previous discussions, respiratory Oscillometry is founded on principles from electrical engineering. Thus, although obtaining respiratory impedance values is relatively straightforward, interpreting these data can be challenging for clinicians [13]. In the diagnostic setting, proper analysis requires specialized training and experience, making it a complex task for professionals not well-versed in the technique [14].

Artificial Intelligence (AI) methods have a long history of contributions to respiratory medicine, dating back to the 1980s [15]. The diagnosis of respiratory disorders has significantly improved with the use of machine learning (ML) techniques in conjunction with oscillometric parameters in asthma [16], small airway dysfunction [17, 18], chronic obstructive pulmonary disease (COPD) [19], sarcoidosis [20], early COPD [18, 21, 22] systemic sclerosis [23] and cystic fibrosis [24]. This technology simplifies and enhances medical decision-making by supplementing human expertise with valuable insights.

Explainable ML methods provide the transparency needed for clinicians to trust AI-assisted diagnoses and treatment recommendations [20]. However, no studies have employed machine learning or explainable methods to enhance the diagnosis of respiratory abnormalities in patients with silicosis.

Therefore, this work uses machine learning and explainable methods to help the medical team investigate and diagnose respiratory alterations in silicosis based on data obtained by oscillometry and respiratory modeling.

This investigation itself presents a few challenges. The first one is the size of the training dataset (sixty volunteers), so we propose the use of synthetic data generators to increase the number of examples that are going to be used for training the machine learning algorithms.

The second one is to employ deep learning algorithms in small datasets. Numerous domains, including computer vision [25], natural language processing [26], video analysis [27], and reinforcement learning [28], have already seen significant success with deep learning. Deep learning techniques are less common in tabular data analysis than in other domains. It turns out that neural networks’ performance on tabular data is inferior to that of other, much simpler methods, even though they were initially designed with this goal in mind [29, 30].

## METHODS

### Ethical issues and training dataset

Prior to commencing this research, ethical approval was obtained from the Medical Research Ethics Committee of the State University of Rio de Janeiro, and registered at ClinicalTrials.gov (identifier: NCT01725971). The studied volunteers were examined from 01/15/2002 to 07/01/2023. This work was developed by collaboration between an R&D laboratory and a teaching tertiary hospital in accordance with the Declaration of Helsinki. Demographic data, including age, height, and weight, were collected from each subject during the procedures. All the volunteers were native South Americans and had to sign a written informed consent form to be included in this study. The research steps are shown in Figure 1.

**Fig. 1:**
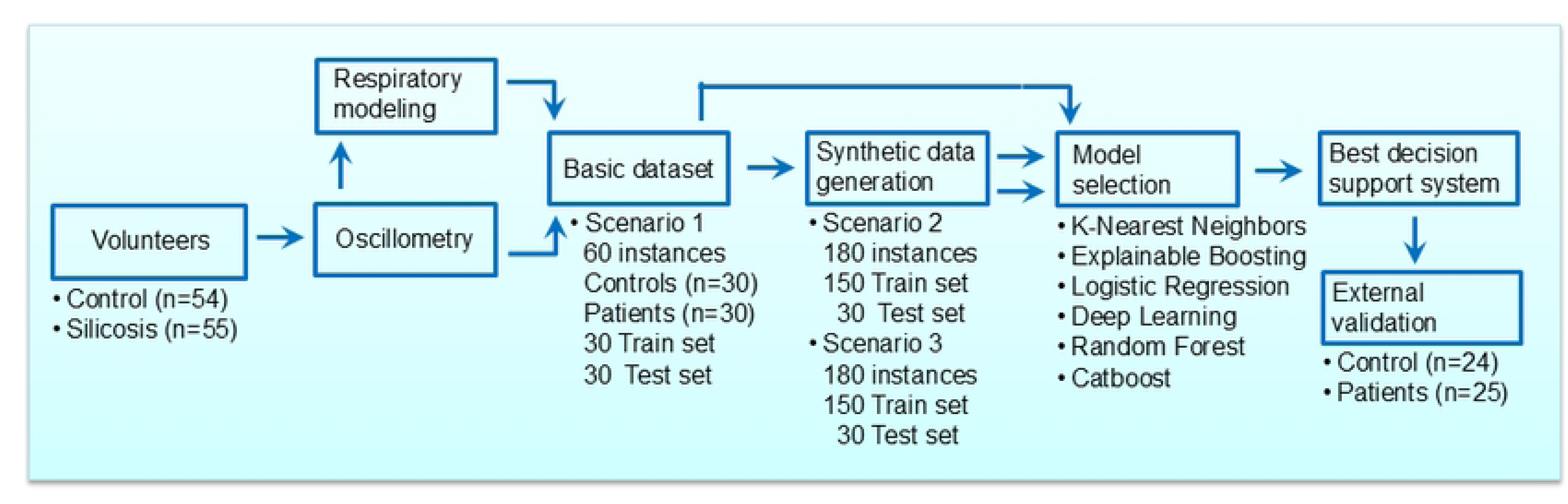
Flowchart of the study.

Individuals without previous lung diseases and with pulmonary function tests compatible with normality were evaluated as a control group. The silicosis group was composed of patients from the outpatient clinic diagnosed according to [31].

In the training group (Figure 1), the exams were conducted on 30 volunteers in the control group and 30 volunteers with Silicosis in the test group, which produced a primary dataset of 60 instances. Exclusion criteria in this group, were history of other lung disease, COVID-19 or smoking.

### Forced oscillation measurements and derived parameters

The dataset used in this work was obtained by an oscillometry system developed in our Laboratory [32], according to standard recommendations [33]. Respiratory impedance (Zrs) is a complex variable calculated as the ratio of the Fourier transform of the small (approximately 2cmH_2_O) excitation pressure (*F*P) and the resulting airflow (*F*V’) at the studied frequencies [Zrs(f)=F(P)/F(V’)].

Three 16-second measurements were performed, and their average was calculated. Tests free from artifacts presenting a coefficient of variation ≤10% among measurements were included [33]. To prevent alterations in bronchial muscle tone caused by the maximum effort required in spirometry, these analyses were conducted before spirometry. Additionally, only tests with a coherence function ≥0.9 across the entire frequency range were accepted to minimize the impact of spontaneous breathing on the measurement of respiratory impedance [34, 35].

Resistive characteristics were evaluated at 4 Hz, 12 Hz and 20 Hz (R4, R12 and R20, respectively), along with the variation between R4 and R20 (R4-R20). Reactance was assessed through several parameters: the dynamic compliance (Cdyn), the resonance frequency (Fr), the area under the reactance curve (Ax), and the impedance modulus at 4 Hz (Z4). Cdyn, which reflects the respiratory compliance, was calculated from the reactance at 4 Hz (Cdyn = 1/2πfX4). The resonance frequency, defined as the point where respiratory reactance becomes zero, indicates the respiratory system’s homogeneity. Ax was determined by the area under the curve between the lowest frequency (4 Hz), its corresponding reactance (X4), and Fr. The respiratory system mechanical load was assessed using the modulus at 4 Hz (Z4), as it encompasses the respiratory load in resistive and elastic characteristics.

### Electrical Model of the Respiratory System

Figure 2 shows a simplified electric model of the oscillometric instrument and the extended resistance-inertance-compliance model (eRIC), which interprets the respiratory system as two-compartment subsystems (central and peripheral) in series [8]. Resistance (R), inductance (I), and capacitance (C) are analogs of respiratory resistance, inertance, and compliance, respectively, while Rp represents the peripheral resistance.

**Fig. 2:**
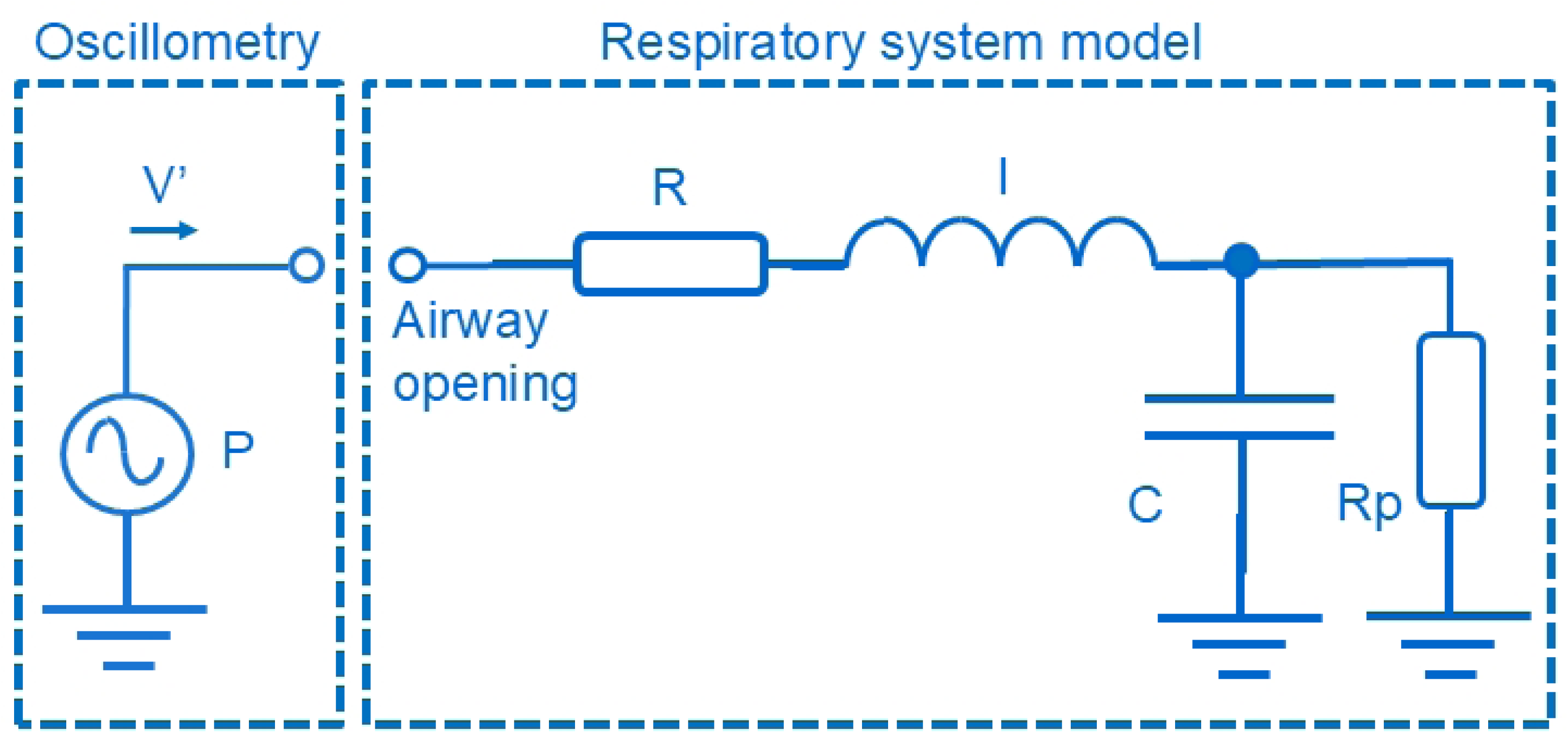
Simplified electric model of the oscillometric instrument describing the excitation pressure (P) and resulting airflow (V’), and the two-compartment (central and peripheral) integer-order model (eRIC) used to analyze respiratory impedance. Resistance (R), inductance (I), and capacitance (C) are analogs of respiratory resistance, inertance, and compliance. Rp represents the peripheral resistance.

Model parameters were estimated within the LABVIEW™ 2018 environment using ModeLIB, a program developed at LIB/UERJ. A Levenberg-Marquardt algorithm was used in this process, determining the nonlinear model coefficients that yielded the best least-squares fit to the input data.

### Machine Learning Algorithms

A subfield of AI, machine learning enables computers to learn without explicit instruction [36]. Its methods work best with problems that lack a deterministic solution, where data is utilized to enable algorithms to automatically determine the link between variables. Prior studies have demonstrated the value of combining oscillometric features with machine learning methods to investigate and support the diagnosis of respiratory illnesses [37–39]. Extensive model testing showed that ensemble methods yielded outstanding results. In the present study, we conduct more experiments and further investigate two distinct ensembles: the Explainable Boosting Machine (EBM) and the Catboost (CAT). The following ML algorithms were also evaluated: Logistic Regression (LR), K-Nearest Neighbor (KNN), Random Forest (RF) [40], and HyperTab, a deep learning approach to small tabular datasets. The first three algorithms have already been briefly detailed in our group’s past investigations [14, 38]. In the following, we present a concise overview of the two algorithms not included in our previous work. Full details can be found in the cited references.

Categorical Boosting (CatBoost) [41] is another implementation of gradient boosting that uses binary decision trees as base predictors. It presents similar and sometimes superior speed and accuracy to algorithms in the same class, such as XGBoost [42] and LightGBM [43]. One of the main differences between CatBoost and other algorithms is its implementation of symmetric trees (or oblivious trees). Oblivious means that the same splitting criteria are used throughout the tree level. These trees are balanced, less prone to overfitting, and significantly speed up model execution at test time. Also, it performs gradient boosting differently, using ordered boosting. CatBoost randomly creates an artificial timestamp for each data point if the data does not have a timestamp. It calculates the residual for each data point using what was trained with the other data points up to that point. Thus, different models are trained to calculate the next residuals for the other data points with data that the model has yet to see. CatBoost divides the dataset into random permits and applies ordered boosting to it. By default, the algorithm creates four permutations. With this randomness, it is possible to avoid further overfitting the model.

As proposed by [44], the Explainable Boosting Machine (EBM) is a glass-box model that preserves intrinsic interpretability while utilizing cutting-edge performing algorithms, such as boosting and bagging. This model’s main goal is to construct a Generalized Additive Model with interactions (GA^2^M) by applying a low learning rate to a round-robin training approach on one feature at a time. This method outperforms the original GAM in two significant ways. First, it can record the significance of feature interactions or the combined effect of two relevant features on the model. Its general is given in the form of (Eq. 1):

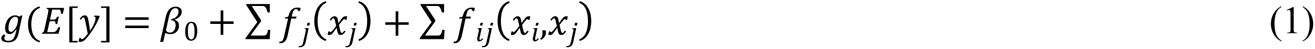

Where *g* is the link function, *f_j_* and *f_ij_*are called smooth functions. This approach brings two significant improvements compared to the original GAM (Hastie; Tibshirani, 1986). Initially, it can document the importance of feature interactions or the joint impact of two model-relevant features. In the context of respiratory physiology, feature interactions are ubiquitous. For example, changes in airway resistance directly impact lung volume. This condition is often disregarded because most explainability algorithms rely on local linear approximations. Retaining interpretability while increasing the model’s accuracy is made possible by capturing the interaction between two features.

The second improvement is FAST, a powerful tool for determining and categorizing the level of interaction between each pair of variables. EBM is a condensed version of the model presented by Yin and his collaborators [45]. It is straightforward to evaluate the relative contributions of individual features and pairwise interactions thanks to the additive structure of this model. As a result, local and global interpretations can be obtained using this method. By computing the result of each smooth function given the feature values, we can interpret the local behavior of the model and understand the influence of individual physiological parameters. Similar to logistic regression feature weights, the mean of smooth functions provides a global interpretation.

A hypernetwork-based method called HyperTab addresses the issue of classifying tiny tabular datasets [46]. HyperTab builds an ensemble of target networks by leveraging a hypernetwork. A hypernetwork is a neural network that produces the target network’s weights, another neural network. Every target network has been designed to process a particular lower-dimensional data view. To obtain lower-dimensional perspectives, a subset of features from the original data is chosen randomly, known as data augmentation. Compared to other tabular data analysis techniques, HyperTab offers several advantages. First, tiny datasets with a far smaller sample size than features—can be handled by HyperTab with ease. This procedure allows HyperTab to effectively perform a non-domain-specific tabular data augmentation by increasing the quantity and diversity of the training data by feature subsetting, which yields many perspectives of each sample.

Secondly, due to its independence from specific attributes, HyperTab can be very robust. Since it is an ensemble where different target networks are trained with different sets of attributes, it is very robust since it does not compromise its performance if a problematic attribute exists. HyperTab can move forward without requiring any additional tuning if target networks that rely on this particular attribute are simply removed.

### Synthetic Data Generation

Synthetic data can thereafter be used to supplement, enhance, and occasionally even replace real data when training ML models. It also removes the risk of exposure that comes with sharing data, allowing ML and other software systems that depend on data to be tested. Several methods exist for acquiring synthetic data. Two methods were used in this work: the first one creates the synthetic data using a machine learning model [47]. The correlation between the original data columns is maintained using this strategy. Additionally, it will respect certain statistical parameters like mean, variance, maximum, and lowest values.

The generative adversarial network (GAN) is the foundation of the second technique. This architecture shows two neural networks: a discriminator network and a generator network. While the latter is trained to determine whether the data is artificially generated, the former is trained to generate artificial data. The rivalry between these networks enables the GAN to mimic any data distribution [48].

It is important to note that the synthetic data was used solely to augment the size of the training data set. This practical approach significantly influenced the results obtained in the test set.

### Experimental design

Our study comprised three experiments. The first experiment evaluated the accuracy of each oscillometric parameter to correctly detect respiratory changes associated with silicosis.

All eleven original oscillometric and modeling parameters were applied to ML algorithms in the second experiment. Four of the six chosen classifiers were implemented with Scikit-learn, a library written in Python. The other two classifiers were also implemented in Python in Hypertab libraries [46] and InterpretML [44]. Since AUC is one of the more widely used metrics in medicine [49] and provides a better method of confronting classifiers than accuracy [50], it was selected as the performance measure. These values were classified according to Greiner et al [51] (0.70<AUC<0.90, moderate; AUC>0.90, high accuracy).

We split the dataset into training and test sets using the hold-out procedure before any preprocessing of the inputs. Each contains thirty measurements to assess how well machine learning models handle tiny datasets. Hyperparameter tuning is a crucial step in model selection, and we applied cross-validation (CV) only to the training data. The Optuna library [52] presents a more efficient strategy to allow hyperparameter fine-tuning while at the same time providing the same interface as gridsearchCV, which is a popular hyperparameter tuning strategy used in the Scikit-learn library [53] with all possible combinations of the hyperparameters. Table 1 presents the classifiers and their respective chosen hyperparameters for tuning.

**Table 1.**
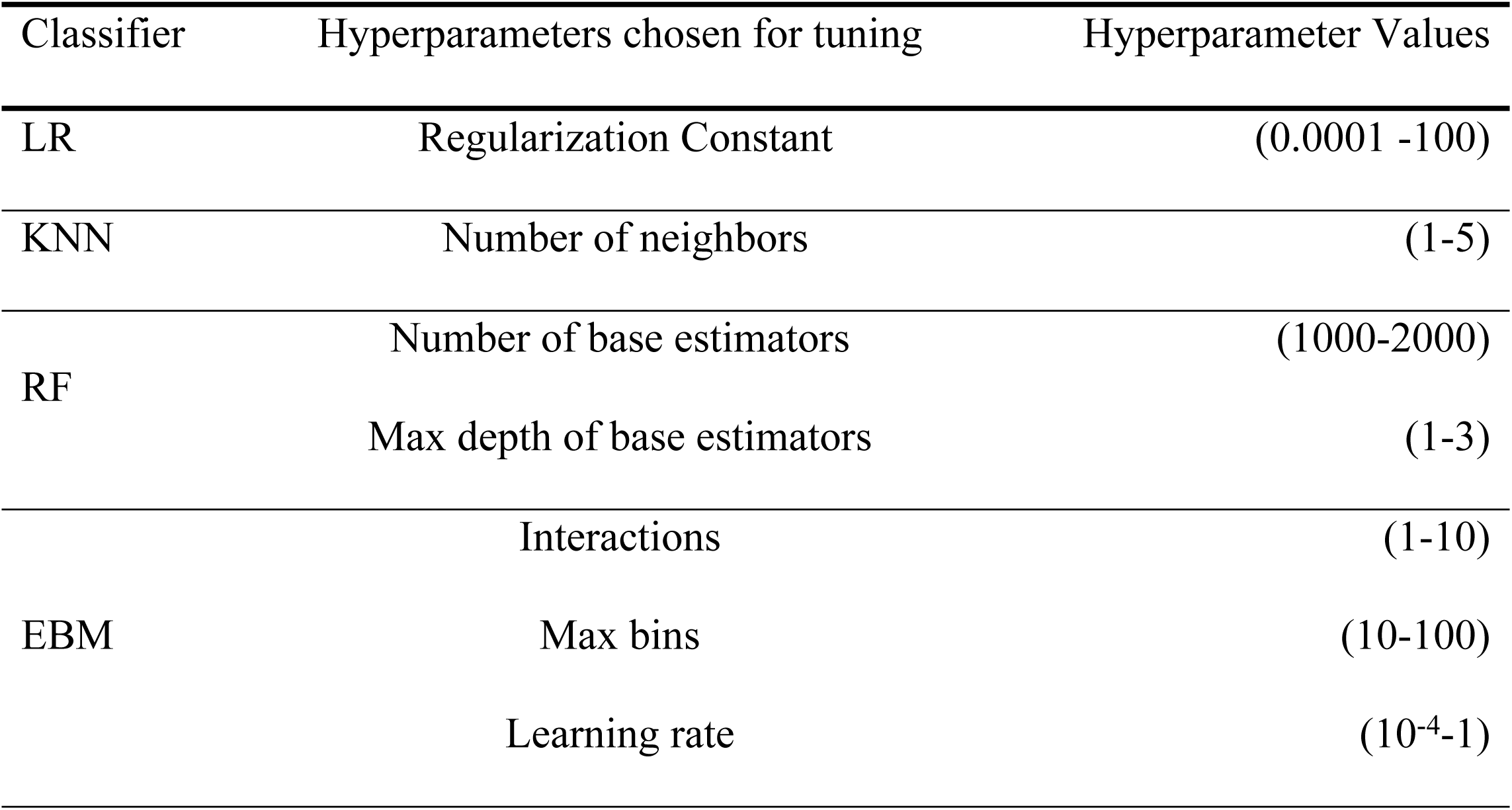
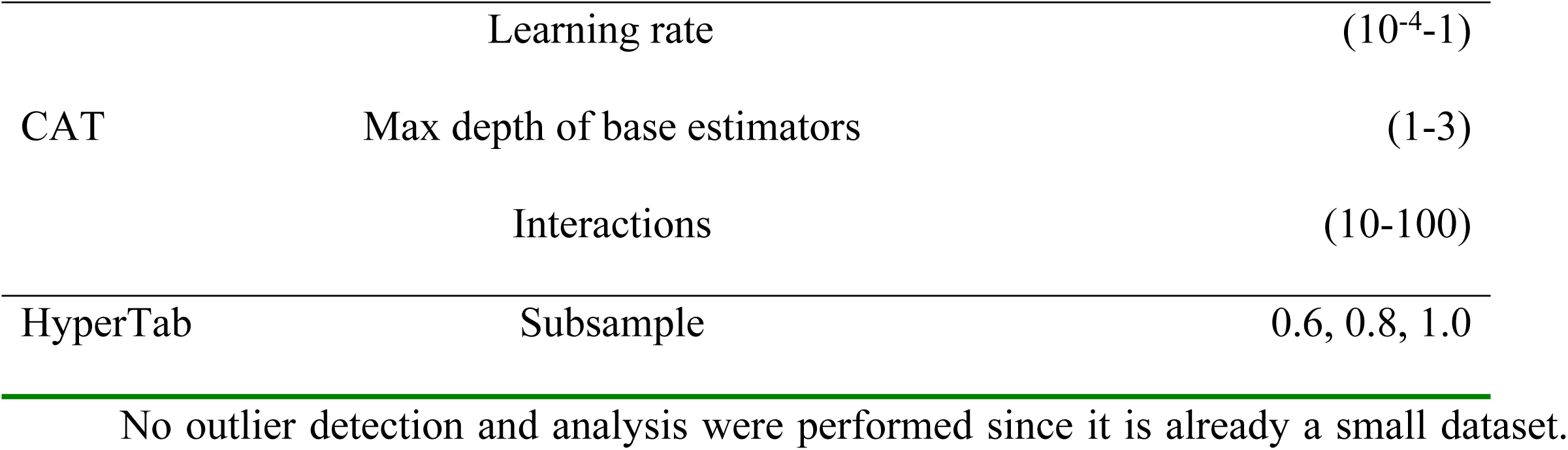
Hyperparameter values used for tuning each used classifier.

No outlier detection and analysis were performed since it is already a small dataset. Also, there was no need for missing value management, the feature preprocessing employed is the standardization, and no data imbalance analysis was performed because the dataset was balanced.

In the third experiment, we aimed to improve algorithm performance by selecting a reduced subset of the initial oscillometric parameters. Based on a wrapper strategy, which identifies input features maximizing average AUC, we employed a Logistic Regressor Classifier for feature selection. To mitigate potential overfitting from this process, and given the small dataset size, two-fold cross-validation was used.

The BFP from the first experiment was then compared against the six other classifiers applied in the second and third experiments.

### External validation

The system was externally validated using a completely independent data set (Figure 1). Our University Hospital receives patients from all over Rio de Janeiro, and patients with silicosis come from very different activities such as mining, construction, quarrying and stone crushing, foundry work, sandblasting, and glass and ceramics manufacturing. Therefore, this new data comes from a different source than the training data, ensuring a true test of generalization. We used the best model in another dataset composed of 24 individuals in the control group and 25 patients with silicosis in the test group. These groups followed the same acceptability criteria used in the training groups.

## RESULTS

### Studied volunteers in the training set

There were no significant differences between the analyzed and the control groups regarding the anthropometric parameters (Table 2). A significant reduction in all spirometric parameters was observed in patients with silicosis.

**Table 2.** Demographics and spirometric characteristics of the subjects in the training set.

|  | Control<br>(n = 30) | Silicosis<br>(n = 30) | p |
| --- | --- | --- | --- |
| Age (years) | 51.3 ± 15.9 | 50.4 ± 13.8 | ns |
| Height (cm) | 171.5 ± 7.9 | 168.7 ± 8.5 | ns |
| Weight (kg) | 74.8 ± 11.8 | 70.0 ± 13.9 | ns |
| BMI (kg/m<sup>2</sup>) | 25.4 ± 3.6 | 24.6 ± 4.3 | ns |
| Gender (M/F) | (22/8) | (23/7) | - |
| Exposition time (years) | - | 15.4 ± 6.3 | - |
| Beginning of symptoms | - | 8.6 ± 8.5 | - |
| Smoking (pack-years) | 0 | 0 | - |
| FEV<sub>1</sub> (L) | 3.5 ± 0.6 | 2.3 ± 0.8 | <0.0001 |
| FEV<sub>1</sub> (%) | 101.8 ± 12.0 | 70.0 ± 20.0 | <0.0001 |
| FVC (L) | 4.3 ± 0.8 | 3.5 ± 1.1 | <0.001 |
| FVC (%) | 101.6 ± 12.5 | 84.8 ± 21.0 | <0.002 |
| FEV<sub>1</sub>/FVC (%) | 90.0 ± 13.4 | 67.6 ± 11.6 | <0.0001 |
| FEF<sub>25-75</sub> (L) | 3.8 ± 1.1 | 1.7 ± 0.7 | <0.0001 |
| FEF<sub>25-75</sub> (%) | 109.8 ± 33.2 | 48.0 ± 20.1 | <0.0001 |

BMI - Body mass index; FEV<sub>1</sub> - Forced expiratory volume in 1 second; FVC - Forced vital capacity; FEV<sub>1</sub>/FVC - Tiffeneau index; FEF<sub>25-75</sub> – Forced expiratory flow between 25% e 75%; n = number of evaluated patients; ns - not significant; (%) - percentile of the predicted values.

### Forced oscillation parameters

Figure 3 displays the resistance and reactance curves obtained from the respiratory oscillometry measurements. The box plots seen in Figure 4 show an increase in the mean values of fr, Z4, R20, R4-R20, R12, Ax, R12, R, Rp, Rt) in patients with silicosis in comparison with the control group. The test group showed decreased mean values for Cdyn, C, and I compared to the control group.

**Fig. 3:**
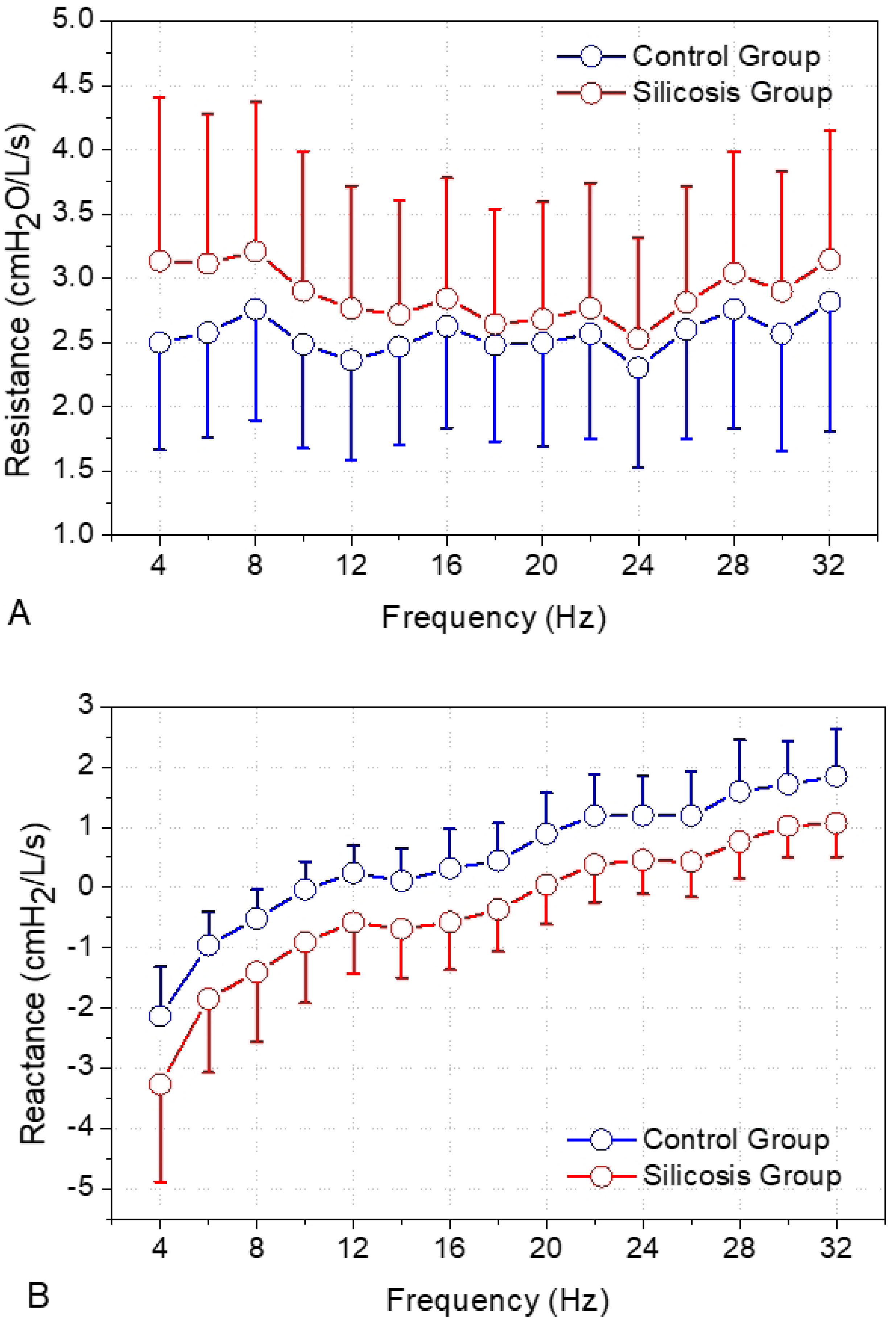
Resistance and reactance curves as a function of frequency obtained from the respiratory oscillometry measurements.

**Fig. 4:**
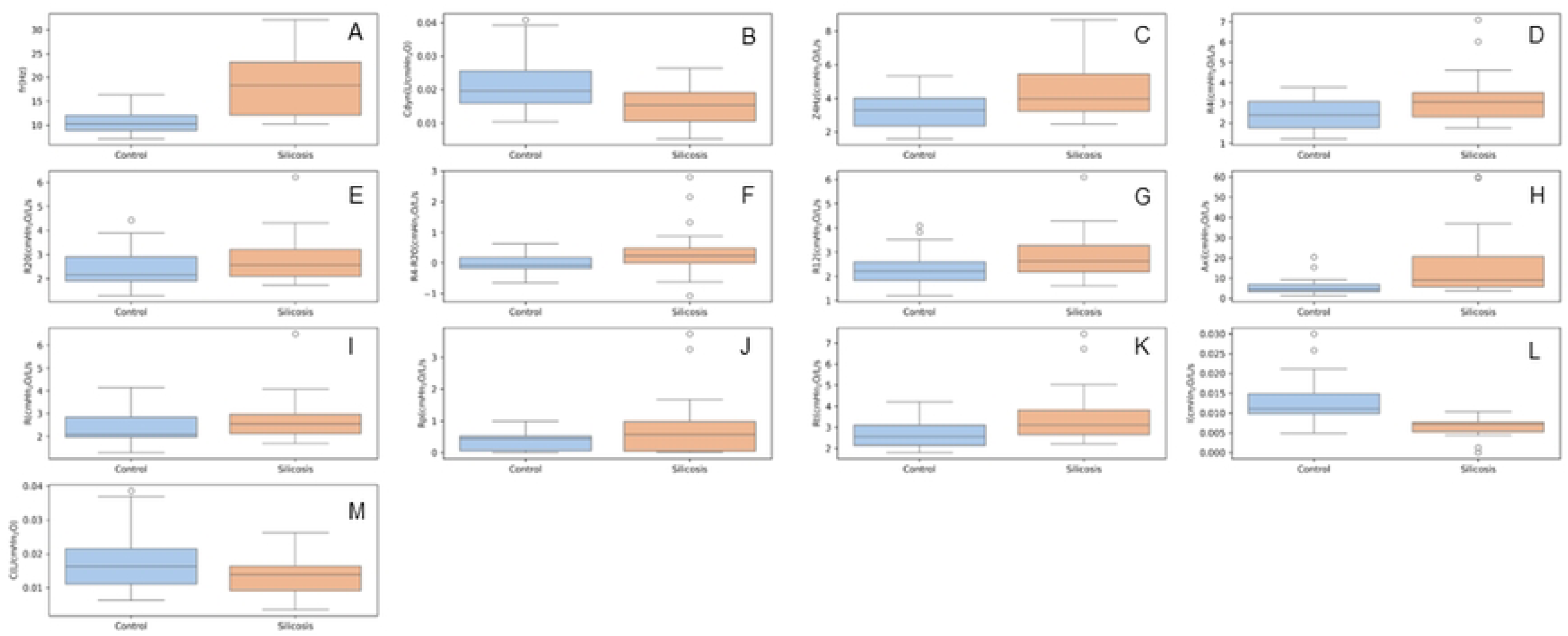
Box plots describing values from the test group compared to the control group in the training set: Resonant frequency (fr, A), dynamic compliance (Cdyn, B), respiratory impedance modulus (Z4, C), resistance in 4 Hz (R4, D), 20 Hz, (R20, E), the difference between R4 and R20 (R4-R20, F), resistance in 12 Hz (R12, G) and area under the reactance curve (Ax, H). Parameters obtained from the electric model are also showed: central resistance (R, I), peripheral resistance (Rp, J), total resistance (Rt, K), respiratory inertance (I, L) and compliance (C, M).

### Experiment 1: Evaluating the diagnostic accuracy of individual FOT parameters

Figure 5 summarizes the results of the first experiment, showing moderate diagnostic accuracy for all parameters (0.70≤AUC≤0.90). The oscillometric parameters f_r_ and I showed the best performance (AUC=0.83).

**Fig. 5:**
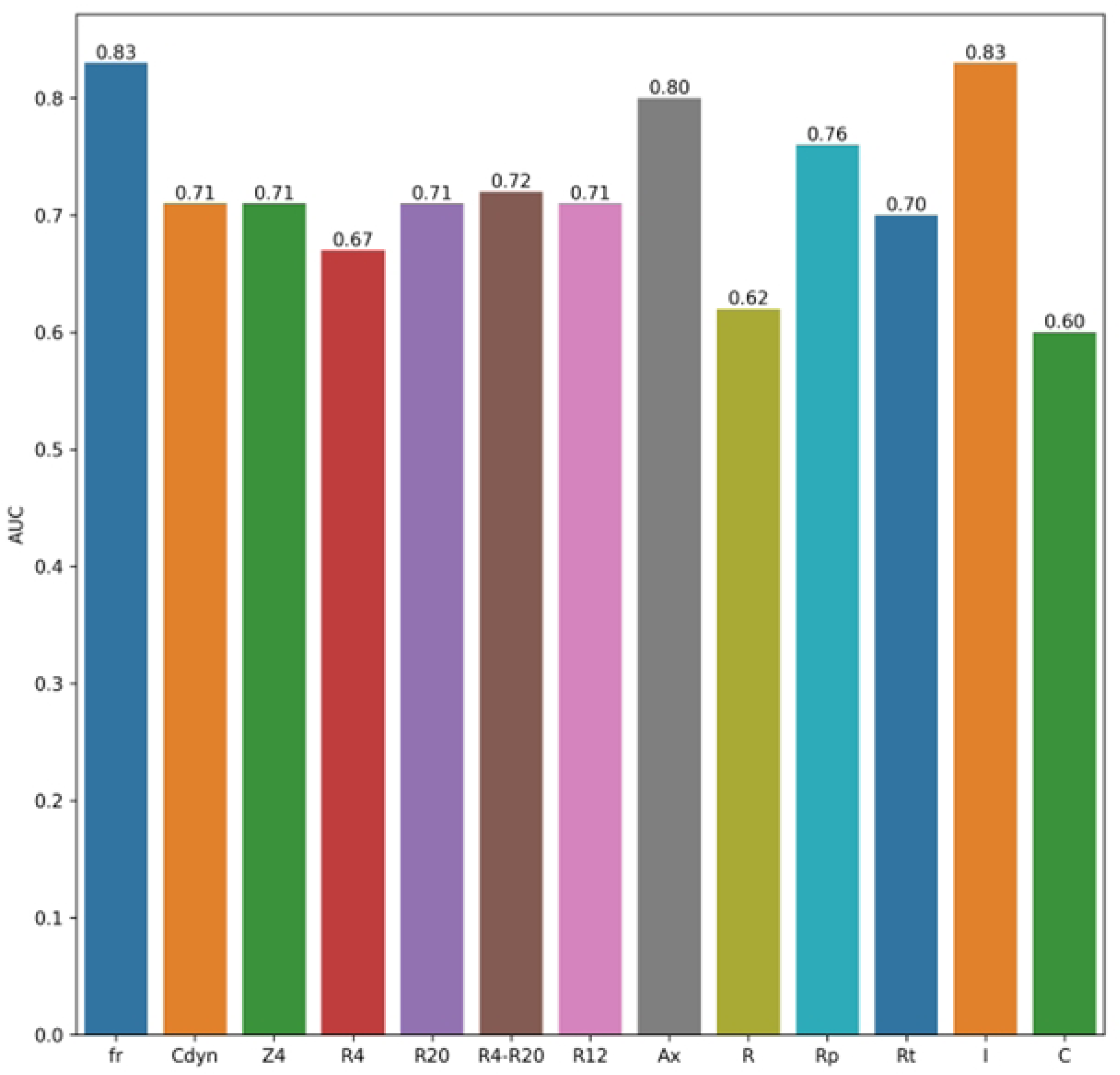
Diagnostic accuracy values obtained in this first experiment, considering only the oscillometric parameters.

The supplementary material includes the ROC curves and tables with the AUC, standard error, sensitivity and specificity and the confidence intervals of each FOT parameter (Figure S1).

### Experiment 2: Evaluating the diagnostic accuracy of FOT combined with Machine Learning

Figure 6 shows the AUC for each classifier and the BFP obtained in this experiment, considering the three different situations to generate synthetic data.

**Fig. 6:**
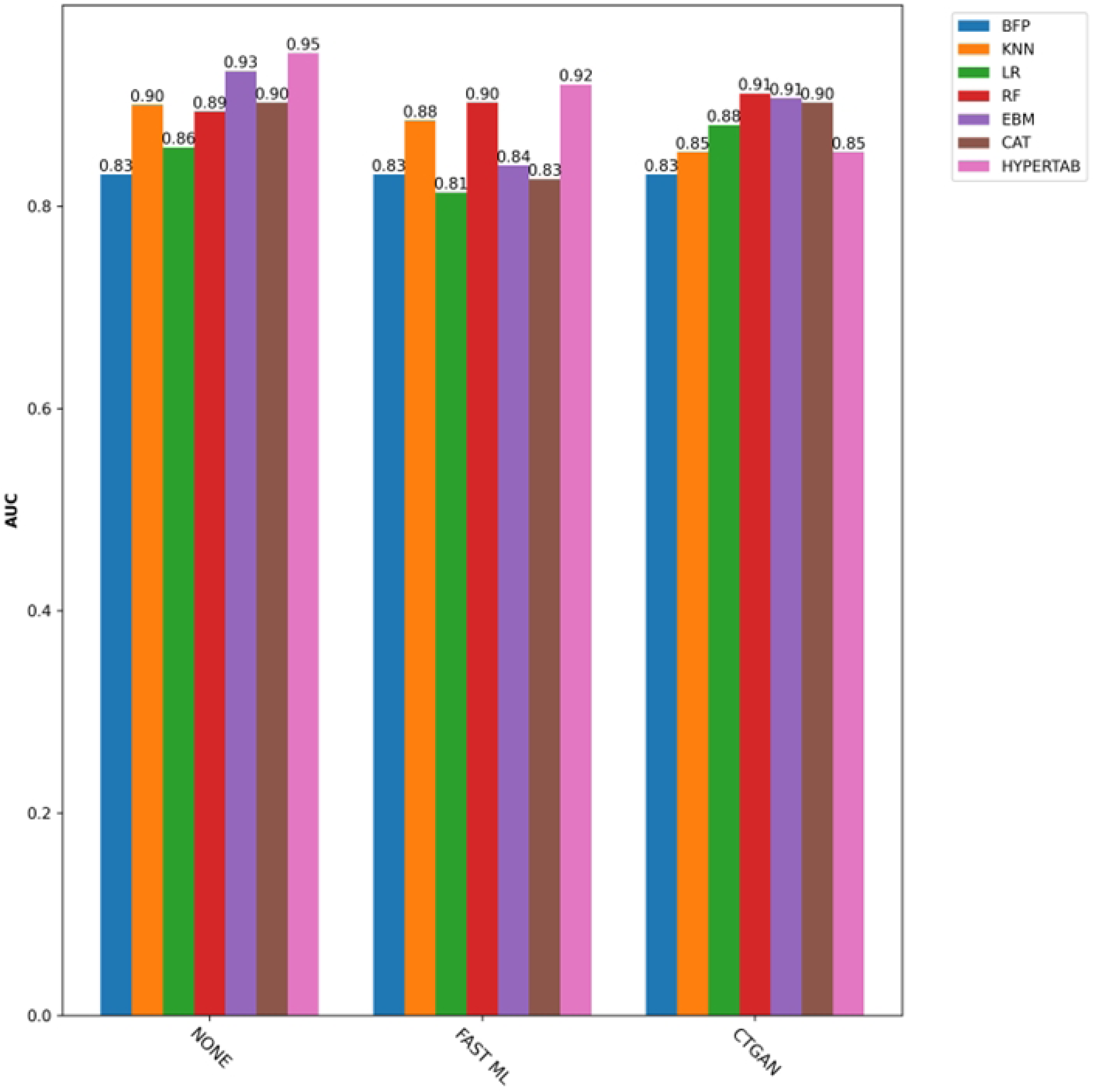
Diagnostic accuracy values obtained for each classifier, the best FOT parameter (BFP), and considering the three different situations used to generate synthetic data.

NONE indicates that no synthetic data was generated, FAST ML describes that data was generated with a machine learning model, and CTGAN signifies that the data was generated with GAN. These two methods are a way to limit the over-fitting due to the sample size. They are performed only on the training data, boosting the training data size to 150 samples. HyperTab and EBM, the top-performing models for the three synthetic data scenarios, achieved excellent AUCs of 0.95 and 0.93, respectively. This success was further highlighted by KNN and CAT, which also achieved an AUC ≥ 0.9, when no synthetic data was generated. In the FAST ML scenario, HyperTab and RF excelled with the best performance, achieving an AUC equal to 0.92 and 0.90, respectively. In the CTGAN scenario, EBM, CAT, and RF were the top performers with AUC ≥ 0.9, further reinforcing the experiment’s success.

### Experiment 3: Evaluating the diagnostic accuracy of the best FOT parameters combined with Machine Learning

After feature selection, the AUCs of the best-performing FOT parameter (Fr) and six other classifiers are shown in Figure 7 (K-NN, LR, RF, EBM, CAT, and HyperTab). EBM and HyperTab outperformed other models, achieving the highest overall AUC (0.94). Considering the NONE case, KNN, EBM, CAT, and RF were the best with AUC ≥ 0.9. In the FAST_ML situation, RF (AUC = 0.90) was the only one that achieved AUC ≥ 0.9. In CTGAN, four classifiers achieved AUC ≥ 0.9: RF, CAT, EBM, and HyperTab.

**Fig. 7:**
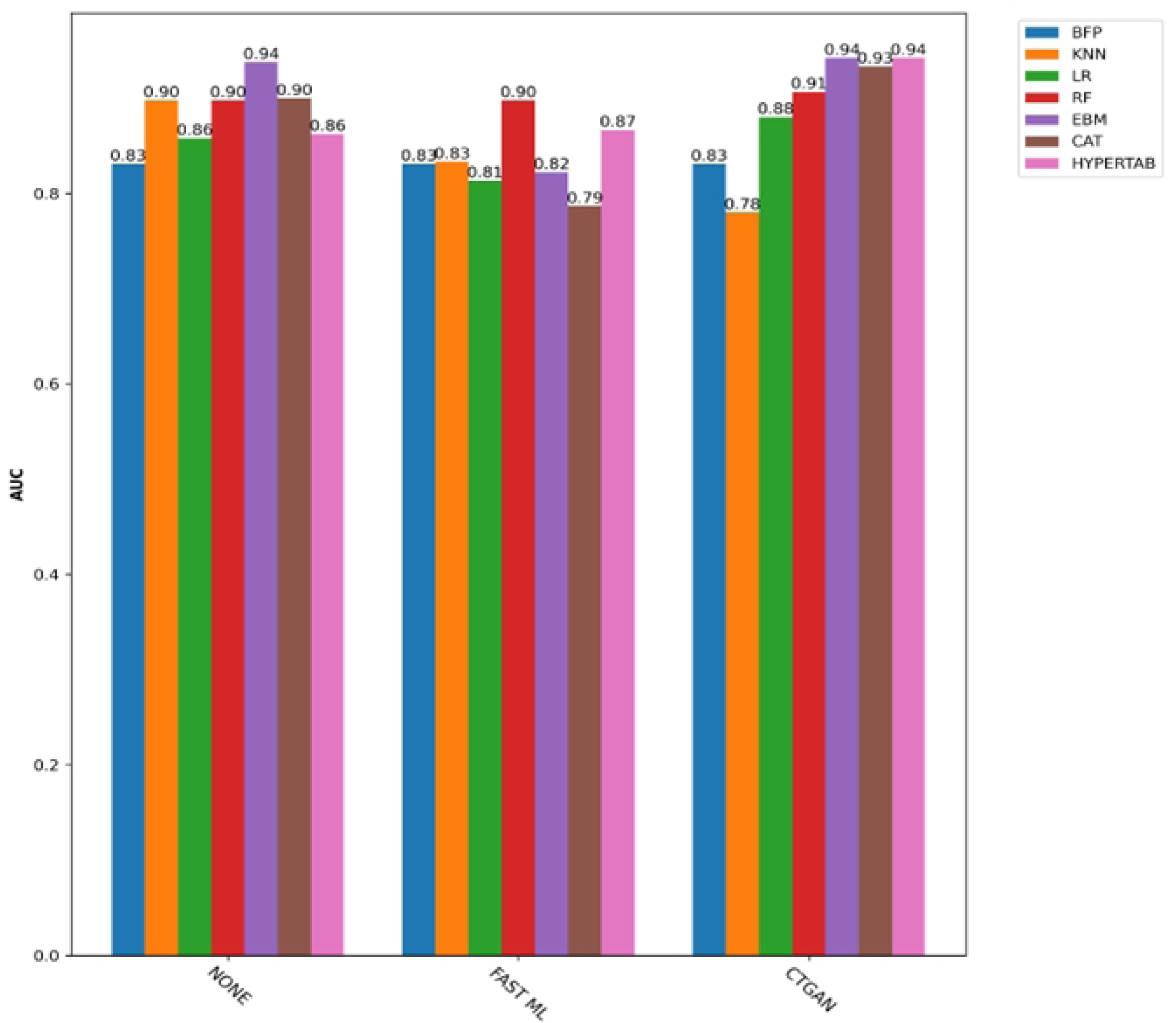
Diagnostic accuracy values obtained using the studied classifiers with feature selection.

Figure 8 shows the importance of features and their interactions, while Figure 9 shows the local explanation for an individual.

**Fig. 8:**
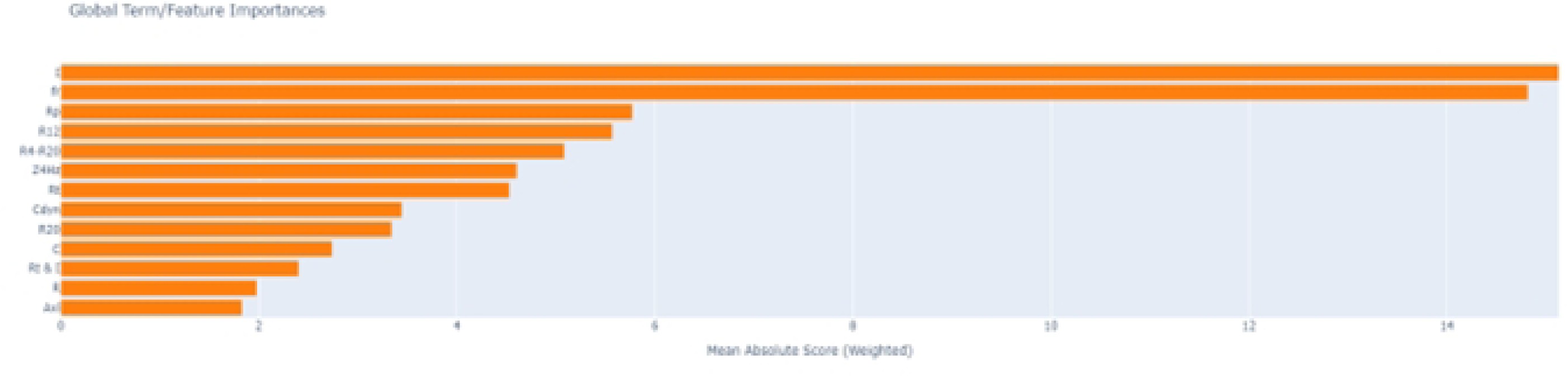
Importance of features and their interactions.

**Fig. 9:**
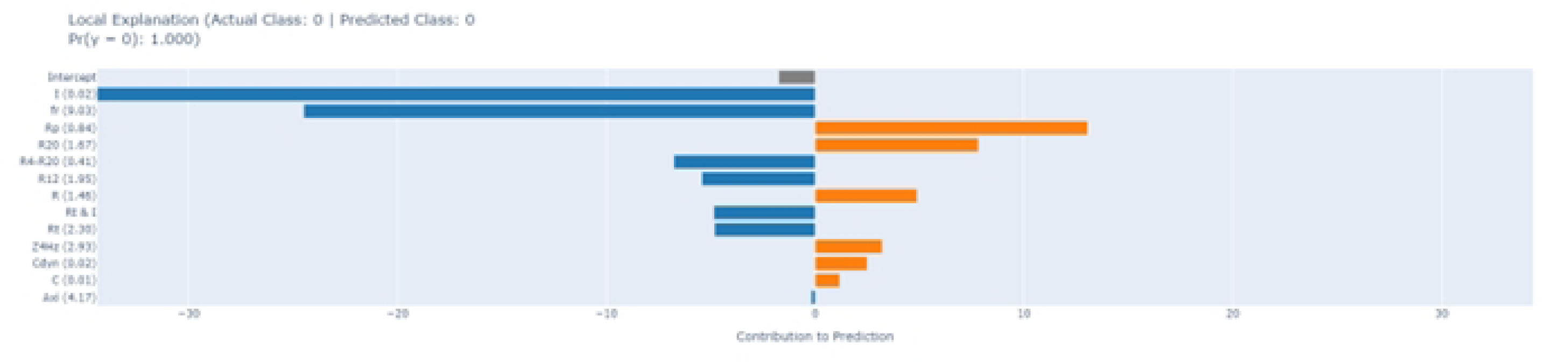
Example of the local explanation for an individual.

Tables 3 and 4 presents Se at a higher Sp (Sp=90%) and Se at a moderate Sp (Sp=75%) in experiments 2 and 3, respectively.

**Table 3.** Sensitivity (%) at 90% Specificity.

|  |  | BFP | KNN | LR | RF | EBM | CAT | HyperTab |
| --- | --- | --- | --- | --- | --- | --- | --- | --- |
| Exp. 2 | NONE | 60.0 | **87.3** | 40.0 | 66.7 | **80.0** | 60.0 | **86.7** |
|  | FAST_ML | 60.0 | **80.8** | 26.7 | 66.7 | 40.0 | 13.3 | 60.0 |
|  | CTGAN | 60.0 | 70.0 | 53.3 | 60.0 | **73.3** | 60.0 | 53.3 |
| Exp 3 | NONE | 60.0 | **73.3** | 40.0 | 60.0 | **80.0** | 60.0 | 40.0 |
|  | FAST_ML | 60.0 | 30.0 | 26.7 | 60.0 | 33.3 | 0.06 | 40.0 |
|  | CTGAN | 60.0 | 15.0 | 53.3 | 53.3 | 66.6 | 66.6 | 80.0 |

**Table 4.** Sensitivity (%) at 75% Sp.

|  |  | BFP | KNN | LR | RF | EBM | CAT | HyperTab |
| --- | --- | --- | --- | --- | --- | --- | --- | --- |
| Exp. 2 | NONE | 66.7 | **90.3** | 73.3 | 86.7 | **93.3** | 80.0 | **100.0** |
|  | FAST_ML | 66.7 | 84.6 | 73.3 | 80.0 | 66.6 | 66.6 | **100.0** |
|  | CTGAN | 66.7 | 81.1 | 73.3 | **93.3** | 86.7 | **93.3** | 60.0 |
| Exp 3 | NONE | 66.7 | 75.8 | 73.3 | 86.7 | 86.7 | 80.0 | 73.3 |
|  | FAST_ML | 66.7 | 80.0 | 73.3 | 66.7 | 80.0 | 66.7 | 73.3 |
|  | CTGAN | 66.7 | 71.7 | 73.3 | **93.3** | **100.0** | **93.3** | 80.0 |

### External validation

Table 5 shows the biometric and spirometric characteristics of the validation set. Using the trained EBM model in the validation set achieved an AUC = 0.96. This value is slightly higher than obtained using the test set in Experiment 2 (AUC = 0.93) and Experiment 3 (AUC = 0.94).

**Table 5.** Subject demographics and spirometric characteristics in the validation set.

|  | Control<br>(n = 24) | Silicosis<br>(n = 25) | p |
| --- | --- | --- | --- |
| Age (years) | 46.9 ± 10.6 | 47.4 ± 12.2 | ns |
| Height (cm) | 164.4 ± 8.0 | 165.2 ± 7.7 | ns |
| Weight (kg) | 69.7 ± 13.2 | 64.9 ± 10.9 | ns |
| BMI (kg/m<sup>2</sup>) | 25.6 ± 3.6 | 23.8 ± 3.7 | ns |
| Gender (M/F) | (11/13) | (23/2) | - |
| Smoking (pack-years) | 0 | 0 | - |
| FEV<sub>1</sub> (L) | 3.1 ± 0.7 | 2.0 ± 0.9 | <0.0001 |
| FEV<sub>1</sub> (%) | 104.8 ± 19.0 | 65.9 ± 26.5 | <0.0001 |
| FVC (L) | 3.8 ± 0.9 | 3.1 ± 1.2 | <0.05 |
| FVC (%) | 103.1 ± 17.4 | 82.2 ± 29.0 | <0.01 |
| FEV<sub>1</sub>/FVC (%) | 101.5 ± 10.1 | 81.1 ± 17.3 | <0.0001 |
| FEF<sub>25-75</sub> (L) | 3.3 ± 1.0 | 1.5 ± 0.9 | <0.0001 |
| FEF<sub>25-75</sub> (%) | 108.1 ± 34.9 | 45.0 ± 29.3 | <0.0001 |

## Discussion

Four principal findings emerged from this study: 1) initially, the use of machine learning models in conjunction with oscillometry and electrical modeling data may achieve an accurate, noninvasive diagnosis of respiratory abnormalities in silicosis; 2) automatic classifiers could increase silicosis diagnosis accuracy; 3) EBM provides clinically important explanations about the model’s diagnosis and 4) external validation confirmed the high accuracy obtained using the training set.

Table 2 shows that the studied groups were homogeneous in the training set, without significant biometric differences. All comparisons of the pulmonary function parameters presented reduced values in patients with silicosis, which is in close agreement with the pathophysiology of silicosis [1].

The findings in Figures 3 and 4 are consistent with previous oscillometric analyses [54]. They are also consistent with the structural relationships observed in patients with silicosis between the computed tomography densitometry values and oscillometric parameters [55]. A similar association between eRIC model parameters and structural abnormalities [56] may explain the observed increases in R, Rp, and Rt (Figures 4I, 4J, and 4K, respectively) and the decreases in I (Figure 4L) and Cdyn (Figure 4M).

In the first experiment, Fr was the FOT parameter that presented the best individual performance (Figure 5, AUC=0.83), achieving a moderate accuracy.

Experiment 2 utilized all FOT parameters as attributes (Figure 6). HyperTab achieved the highest overall performance with an AUC of 0.95, while KNN, RF, EBM, and CAT also demonstrated high performance with AUCs of 0.9 or greater. KNN, EBM, and HyperTab maintained moderate sensitivity (70-90%) at a specificity of 90% (Table 3).

Experiment 3 (Figure 7) used the best FOT parameters as input in all studied classifiers. This choice of parameters coincided with the feature selection conducted by recursive feature elimination (RFE) employing logistic regression as a base classifier. Six parameters were selected: Fr, R4-R20, Ax, Rp, Rt, I. The EBM and HyperTab algorithms had the highest performance (AUC=0.94). Considering the synthetic data generation, CTGAN was the most useful since it enhanced the ML models’ performance. In Experiment 3, KNN and EBM achieved moderate Se. Regarding 75% specificity (Table 4), KNN, RF, EBM, CAT, and HyperTab obtained high sensitivity (≥ 90%) in experiments 2 and 3, indicating ML methods’ efficacy in detecting respiratory changes in silicosis.

EBM and HyperTab presented the best scores in the experiments (Figures 6 and 7). However, EBM has one advantage over HyperTab: the ability to explain the importance of the features and their interactions. The EBM model considers I the most important feature (Figure 8), followed by fr, Rp, R12, R4-R20, Z4, Rt, Cdyn, R20, C, the interaction Rt, I, R, and Ax. The importance of fr and I found by EBM, agrees with the performance of individual FOT parameters. The importance of Rp, R12, and R4-R20 reflects the difficulty in breathing related to the airwaýs total resistance, especially in the peripheral airway (Rp). The biomechanical interpretation of Z4 relates to the work required to overcome resistive and elastic forces, as reflected in the pressure-volume curve of the lungs. Also, it is interesting to note that five top seven features agree with the best features found by feature selection. Another finding is the importance of the interaction of I with Rt (total resistance). These interactions align with the eRIC model, which has the resistance in series with the inertance and C (compliance). These parameters can be used to define fr and Z4.

EBM is also capable of explaining the model’s decision about an individual. Figure 9 shows the local explanation for an individual in the control group. I and fr have substantial negative contributions, which means they contribute to the model’s decision that this individual does not have respiratory alterations. The EBM model considers I the most important feature, followed by fr, Rp, R12, R4-R20, Z4, Rt, Cdyn, R20, C, the interaction Rt& I, R, and Ax. The importance of fr and I agree with the performance of individual FOT parameters. The importance of Rp, R12, and R4-R20 reflects the difficulty in breathing related to the airwaýs total resistance, especially in the peripheral airway (Rp). The importance of Z4 is related to the respiratory muscles’ work to overthrow the resistive and elastic loads. Also, it is interesting to note that five top seven features agree with the best features found by feature selection. Another finding is the importance of the interaction of I with Rt (total resistance). These interactions concur with the eRIC model, which has the resistance in series with the inertance and with C (compliance). These parameters can be used to define fr and Z4.

We performed external validation using an independent dataset of typical patients to evaluate model generalizability and robustness in real-world clinical applications. During this study stage, the trained EBM model achieved high diagnostic accuracy (AUC>0.90). Interestingly, this accuracy was slightly higher than that obtained in the test set in Experiment 2 (AUC=0.93) and in Experiment 3 (AUC=0.94). Mean values for FEV_1_ (%), FEV (%), and FEF25-75 (%) of 70.0, 84.8, and 48.0, respectively, were observed in the training set (Table 2). In the validation set, these mean values were slightly reduced to 65.9, 82.2, and 45.0, respectively (Table 5), describing the presence of increased abnormalities in this group.

It is important to emphasize that we were careful to use non-smoking patients during the system development phase (Table 2). The presence of this additional abnormal factor would make the diagnosis of respiratory changes simpler. After developing the system in these adverse conditions, we kept the same conditions during external validation and still obtained highly accurate results. In clinical practical conditions, the presence of smoking is common in this class of patients. As smoking introduces an additional factor in terms of respiratory abnormalities [1, 57], it is expected that in these conditions, the performance would be even better. Therefore, in the context of the study’s goals, the results obtained during external validation confirmed the high clinical potential of the proposed system.

The accuracy obtained during external validation was similar to that recently obtained by Wang et al. [58], which obtained AUCs of 0.90 and 0.87 in a system to diagnose respiratory diseases based on computed tomography (CT) and chest X-ray, respectively. It is also similar to that obtained recently by Zhou et al. [59] in a model for automated lung lesion segmentation and early prediction of acute respiratory distress syndrome, in which the AUC reached 0.90.

From the systems identification point of view, spirometric exams may be considered a step excitation method in which the patient produces the signal under analysis. Step responses directly reveal a system’s transient behavior, which is crucial in applications where settling time, overshoot, and stability are critical considerations. In the particular case of spirometry, step excitation offers simplicity and direct insight into transient response, characterized by the forced expiratory volume in one second (VEF_1_), a key parameter in this analysis [60]. However, step responses are more susceptible to noise interference, potentially obscuring the system’s underlying behavior.

A long research and development history has established robust techniques for analyzing system responses to sinusoidal excitation [61]. Accurately identifying frequency response characteristics based on sinusoidal excitation, combined with techniques like Fourier analysis, effectively filters out noise, providing a clearer picture of the system’s true response. Its versatility makes it a valuable tool across various engineering and scientific disciplines, from mechanical and electrical to chemical and biological. In the specific case of the respiratory system, the use of external excitation allows a simple examination allowing the analysis in conditions where traditional pulmonary function is not applicable, including abnormalities in premature babies [62], sleep disorders [63], critically ill patients [64] and during cardiac surgery [64].

However, some significant limitations must be overcome before this important technological innovation can be widely used clinically. First, as noted by recent consensus guidelines [65], diagnostic simplicity is crucial for busy non-specialist clinicians. The complexity of interpreting oscillometric data can delay diagnosis and treatment, particularly in settings with limited access to specialists and time or resources for extensive training. This study describes an accurate diagnostic support system that may improve the medical services offered to patients with silicosis, simplifying the use of oscillometry and improving the diagnostic of respiratory abnormalities in these patients.

A critical discussion of this study’s limitations is warranted. Firstly, although the proposed system had good discrimination in external validation and may be used by other health systems for case-identification, discrepancies in feature representation may arise due to differences in the characteristics of the patient cohorts. Patients were collected from one state in Brazil (Rio de Janeiro), potentially introducing selection bias and limiting the generalizability of our findings to other regions. Interested readers can assess the generalizability of the findings to their patient populations by reviewing the anthropometric descriptions and the inclusion and exclusion criteria outlined in this paper for the training and validation sets.

Secondly, our study was limited by a relatively small sample size, which may impact the generalizability of the findings. Future studies with larger and more diverse cohorts are needed.

Another noteworthy point is that the model’s real-time performance in clinical settings requires further evaluation. Future studies should assess its integration into clinical workflows and its impact on patient outcomes. Despite its preliminary nature, this analysis contributes significantly to the ongoing discussion of machine learning in respiratory diseases [58, 59] by providing novel data for silicosis, a disease currently unexplored with machine learning approaches.

Whole-breath impedance measurements, as used in this study, provide a global assessment of lung mechanics. In contrast, within-breath analysis offers the potential to capture finer-grained, dynamic changes in impedance throughout the respiratory cycle [66]. Future research incorporating within-breath impedance parameters is recommended.

## Conclusions

Using FOT parameters alone, it was demonstrated a limited diagnostic accuracy for detecting respiratory changes in silicosis, with AUC values below 0.9 for all parameters. Achieving high diagnostic accuracy (AUC ≥ 0.9) required combining machine learning classifiers with synthetic data generation algorithms, specifically FAST_ML and CTGAN. CTGAN, particularly when combined with feature selection, yielded superior results. Notably, HyperaTab, a deep learning algorithm designed for small datasets, performed well both with and without synthetic data augmentation. EBM also provides good results (AUC≥0.9). In addition, it gives the importance of the features and their interactions and allows us to investigate further why the model gives a particular result to the subject. External validation achieved high diagnostic accuracy (AUC>0.90). These findings demonstrate the potential of machine learning, combined with oscillometry and electric models, to accelerate medical workflows and improve the diagnosis of respiratory abnormalities in silicosis. Further research and validation are needed to translate this potential into clinical practice.

## References

1. Leung CC, Yu ITS, Chen W. Silicosis. The Lancet. 2012;379(9830):2008-18. doi: 10.1016/S0140-6736(12)60235-9.

2. Minelli G, Zona A, Cavariani F, Comba P, Pasetto R. Silicosis mortality in Italy: temporal trends 1990-2012 and spatial patterns 2000-2012. Annali dell’Istituto Superiore di Sanità. 2017;53(4):275–82.

3. Cox CW, Rose CS, Lynch DA. State of the Art: Imaging of Occupational Lung Disease. Radiology. 2014;270(3):681–96. doi: 10.1148/radiol.13121415.

4. Kaminsky DA, Irvin CG. New insights from lung function. Current Opinion in Allergy and Clinical Immunology. 2001;1(3):205–9. doi: 10.1097/00130832-200106000-00002.

5. King GG, Bates J, Berger KI, Calverley P, de Melo PL, Dellacà RL, et al. Technical standards for respiratory oscillometry. European Respiratory Journal. 2020;55(2).

6. Kaminsky DA, Simpson SJ, Berger KI, Calverley P, De Melo PL, Dandurand R, et al. Clinical significance and applications of oscillometry. European respiratory review. 2022;31(163).

7. Bates JHT. Lung mechanics : an inverse modeling approach. 1st Edition ed. Cambridge: Cambridge University Press; 2009. xvi, 220 p. p.

8. Lima AN, Faria AC, Lopes AJ, Jansen JM, Melo PL. Forced oscillations and respiratory system modeling in adults with cystic fibrosis. Biomedical engineering online. 2015;14:11. Epub 2015/04/19. doi: 10.1186/s12938-015-0007-7. PubMed PMID: 25889005; PubMed Central PMCID: PMCPMC4334397.

9. Faria AC, Veiga J, Lopes AJ, Melo PL. Forced oscillation, integer and fractional-order modeling in asthma. Computer methods and programs in biomedicine. 2016;128:12–26. Epub 2016/04/05. doi: 10.1016/j.cmpb.2016.02.010. PubMed PMID: 27040828.

10. de Sa PM, Castro HA, Lopes AJ, Melo PL. Early Diagnosis of Respiratory Abnormalities in Asbestos-Exposed Workers by the Forced Oscillation Technique. PloS one. 2016;11(9):e0161981. Epub 2016/09/10. doi: 10.1371/journal.pone.0161981. PubMed PMID: 27612198; PubMed Central PMCID: PMCPMC5017649.

11. Ribeiro CO, Faria ACD, Lopes AJ, Melo PL. Early Diagnosis of the Effects of Smoking and Chronic Obstructive Pulmonary Disease based on Forced Oscillations and Fractional-order Modelling XXVI Congresso Brasileiro de Engenharia Biomédica - CBEB 2018; Búzios, Rio de Janeiro: Springer, The International Federation for Medical and Biological Engineering (IFMBE) Proceedings book series.; 2018.

12. Ribeiro CO, Lopes AJ, de Melo PL. Oscillation Mechanics, Integer and Fractional Respiratory Modeling in COPD: Effect of Obstruction Severity. International journal of chronic obstructive pulmonary disease. 2020;15:3273–89. doi: 10.2147/COPD.S276690. PubMed PMID: 33324050; PubMed Central PMCID: PMC7733470.

13. Kostorz-Nosal S, Jastrzębski D, Błach A, Skoczyński S. Window of opportunity for respiratory oscillometry: A review of recent research. Respir Physiol Neurobiol. 2023;316:104135. Epub 2023/08/04. doi: 10.1016/j.resp.2023.104135. PubMed PMID: 37536553.

14. Amaral JLM, Lopes AJ, Jansen JM, Faria ACD, Melo PL. An improved method of early diagnosis of smoking-induced respiratory changes using machine learning algorithms. Computer Methods and Programs in Biomedicine. 2013;112(3):441–54. doi: 10.1016/j.cmpb.2013.08.004.

15. do Amaral JLM, de Melo PL. Clinical decision support systems to improve the diagnosis and management of respiratory diseases. In: Barh D, editor. Artificial Intelligence in Precision Health: Academic Press; 2020. p. 359–91.

16. Amaral JL, Lopes AJ, Veiga J, Faria AC, Melo PL. High-accuracy Detection of Airway Obstruction in Asthma Using Machine Learning Algorithms and Forced Oscillation Measurements Computer methods and programs in biomedicine. 2017;144:113–25. doi: 10.1016/j.cmpb.2017.03.023.

17. Xu WJ, Shang WY, Feng JM, Song XY, Li LY, Xie XP, et al. Machine learning for accurate detection of small airway dysfunction-related respiratory changes: an observational study. Respir Res. 2024;25(1):286. Epub 2024/07/26. doi: 10.1186/s12931-024-02911-1. PubMed PMID: 39048993; PubMed Central PMCID: PMCPMC11270925.

18. Amaral JL, Lopes AJ, Jansen JM, Faria AC, Melo PL. An improved method of early diagnosis of smoking-induced respiratory changes using machine learning algorithms. Computer methods and programs in biomedicine. 2013;112(3):441–54. Epub 2013/09/05. doi: 10.1016/j.cmpb.2013.08.004. PubMed PMID: 24001924.

19. Amaral JL, Faria AC, Lopes AJ, Jansen JM, Melo PL. Automatic identification of Chronic Obstructive Pulmonary Disease Based on forced oscillation measurements and artificial neural networks. Conference proceedings : Annual International Conference of the IEEE Engineering in Medicine and Biology Society IEEE Engineering in Medicine and Biology Society Conference. 2010;2010:1394–7. doi: 10.1109/IEMBS.2010.5626727. PubMed PMID: 21096340.

20. Lima AD, Lopes AJ, Amaral JL, Melo PL. Explainable machine learning and respiratory oscillometry for the diagnosis of respiratory abnormalities in sarcoidosis. Plos One. 2022.

21. Shen X, Liu H. Using machine learning for early detection of chronic obstructive pulmonary disease: a narrative review. Respir Res. 2024;25(1):336. Epub 2024/09/10. doi: 10.1186/s12931-024-02960-6. PubMed PMID: 39252086; PubMed Central PMCID: PMCPMC11385799.

22. Wu F, Zhou Y, Peng J, Deng Z, Wen X, Wang Z, et al. Rationale and design of the Early Chronic Obstructive Pulmonary Disease (ECOPD) study in Guangdong, China: a prospective observational cohort study. J Thorac Dis. 2021;13(12):6924–35. Epub 2022/01/25. doi: 10.21037/jtd-21-1379. PubMed PMID: 35070376; PubMed Central PMCID: PMCPMC8743397.

23. Andrade DSM, Ribeiro LM, Lopes AJ, Amaral JLM, Melo PL. Machine learning associated with respiratory oscillometry: a computer-aided diagnosis system for the detection of respiratory abnormalities in systemic sclerosis. Biomedical engineering online. 2021;20(1):31. doi: 10.1186/s12938-021-00865-9. PubMed PMID: 33766046; PubMed Central PMCID: PMC7995797.

24. Pinto NP, Amaral JLM, Lopes AJ, Melo PL. Diagnosis of Respiratory Changes in Cystic Fibrosis Using a Soft Voting Ensemble with Bayesian Networks and Machine Learning Algorithms. Journal of Medical and Biological Engineering. 2023;43(1):112–23. doi: 10.1007/s40846-023-00777-0.

25. Krizhevsky A, Sutskever I, Hinton GE. ImageNet classification with deep convolutional neural networks. Commun ACM. 2017;60(6):84–90. doi: 10.1145/3065386.

26. Young T, Hazarika D, Poria S, Cambria E. Recent Trends in Deep Learning Based Natural Language Processing. arXiv; 2018.

27. Michelsanti D, Tan Z-H, Zhang S-X, Xu Y, Yu M, Yu D, et al. An Overview of Deep-Learning-Based Audio-Visual Speech Enhancement and Separation. arXiv; 2020.

28. Mnih V, Kavukcuoglu K, Silver D, Graves A, Antonoglou I, Wierstra D, et al. Playing Atari with Deep Reinforcement Learning. arXiv; 2013.

29. Shwartz-Ziv R, Armon A. Tabular data: Deep learning is not all you need. Information Fusion. 2022;81:84–90. doi: 10.1016/j.inffus.2021.11.011.

30. Grinsztajn L, Oyallon E, Varoquaux G. Why do tree-based models still outperform deep learning on tabular data? : arXiv; 2022.

31. ILO ILO. Guidelines for the use of the ILO International Classification of Radiographs of Pneumoconioses Geneva International Labour Organization; 2022.

32. de Melo PL, Werneck MM, Giannella-Neto A. New impedance spectrometer for scientific and clinical studies of the respiratory system. Review of Scientific Instruments. 2000;71(7):2867–72. PubMed PMID: WOS:000087906800041.

33. King GG, Bates J, Berger KI, Calverley P, de Melo PL, Dellaca RL, et al. Technical standards for respiratory oscillometry. The European respiratory journal. 2020;55(2). doi: 10.1183/13993003.00753-2019. PubMed PMID: 31772002.

34. Bates JH, Irvin CG, Farre R, Hantos Z. Oscillation mechanics of the respiratory system. Compr Physiol. 2011;1(3):1233–72. doi: 10.1002/cphy.c100058. PubMed PMID: 23733641.

35. Teixeira EM, Ribeiro CO, Lopes AJ, de Melo PL. Respiratory Oscillometry and Functional Performance in Different COPD Phenotypes. Int J Chron Obstruct Pulmon Dis. 2024;19:667–82. Epub 2024/03/11. doi: 10.2147/copd.S446085. PubMed PMID: 38464561; PubMed Central PMCID: PMCPMC10924760.

36. El Naqa I, Murphy MJ. What is machine learning?: Springer; 2015 2015.

37. Amaral JLM, Lopes AJ, Veiga J, Faria ACD, Melo PL. High-accuracy detection of airway obstruction in asthma using machine learning algorithms and forced oscillation measurements. Computer Methods and Programs in Biomedicine. 2017;144:113–25. doi: 10.1016/j.cmpb.2017.03.023.

38. Amaral JLM, Lopes AJ, Jansen JM, Faria ACD, Melo PL. Machine learning algorithms and forced oscillation measurements applied to the automatic identification of chronic obstructive pulmonary disease. Computer methods and programs in biomedicine. 2012;105(3):183–93.

39. Andrade DSM, Ribeiro LM, Lopes AJ, Amaral JLM, Melo PL. Machine learning associated with respiratory oscillometry: a computer-aided diagnosis system for the detection of respiratory abnormalities in systemic sclerosis. BioMed Eng OnLine. 2021;20(1):31. doi: 10.1186/s12938-021-00865-9.

40. Hastie T, Tibshirani R, Friedman JH, Friedman JH. The elements of statistical learning: data mining, inference, and prediction: Springer; 2009 2009.

41. Dorogush AV, Ershov V, Gulin A. CatBoost: gradient boosting with categorical features support. arXiv; 2018.

42. Chen T, He T, Benesty M, Khotilovich V, Tang Y, Cho H, et al. Xgboost: extreme gradient boosting. R package version 04-2. 2015;1(4):1–4.

43. Ke G, Meng Q, Finley T, Wang T, Chen W, Ma W, et al. Lightgbm: A highly efficient gradient boosting decision tree. Advances in neural information processing systems. 2017;30.

44. Nori H, Jenkins S, Koch P, Caruana R. InterpretML: A Unified Framework for Machine Learning Interpretability. arXiv; 2019.

45. Lou Y, Caruana R, Gehrke J, Hooker G, editors. Accurate intelligible models with pairwise interactions. KDD’ 13: The 19th ACM SIGKDD International Conference on Knowledge Discovery and Data Mining; 2013 2013/08/11/. Chicago Illinois USA: ACM.

46. Wydmański W, Bulenok O, Śmieja M. HyperTab: Hypernetwork Approach for Deep Learning on Small Tabular Datasets. arXiv; 2023.

47. Patki N, Wedge R, Veeramachaneni K, editors. The Synthetic Data Vault. 2016 IEEE International Conference on Data Science and Advanced Analytics (DSAA); 2016 17-19 Oct. 2016.

48. Goodfellow IJ, Pouget-Abadie J, Mirza M, Xu B, Warde-Farley D, Ozair S, et al., editors. Generative Adversarial Nets. Neural Information Processing Systems; 2014.

49. Nahm FS. Receiver operating characteristic curve: overview and practical use for clinicians. Korean J Anesthesiol. 2022;75(1):25–36. doi: 10.4097/kja.21209.

50. Japkowicz N, Shah M. Evaluating learning algorithms: a classification perspective: Cambridge University Press; 2011 2011.

51. Greiner M, Pfeiffer D, Smith RD. Principles and practical application of the receiver-operating characteristic analysis for diagnostic tests. 2000;45(1-2):23–41.

52. Akiba T, Sano S, Yanase T, Ohta T, Koyama M, editors. Optuna: A Next-generation Hyperparameter Optimization Framework2019 2019.

53. Pedregosa F, Varoquaux G, Gramfort A, Michel V, Thirion B, Grisel O, et al. Scikit-learn: Machine Learning in Python. Journal of Machine Learning Research. 2011;12:2825–30.

54. de Sa PM, Lopes AJ, Jansen JM, de Melo PL. Oscillation mechanics of the respiratory system in never-smoking patients with silicosis: pathophysiological study and evaluation of diagnostic accuracy. Clinics (Sao Paulo). 2013;68(5). doi: 10.6061/clinics/2013(05)11. PubMed PMID: 23778400; PubMed Central PMCID: PMC3654297.

55. Lopes AJ, Mogami R, Camilo GB, Machado DC, Melo PL, Carvalho AR. Relationships between the pulmonary densitometry values obtained by CT and the forced oscillation technique parameters in patients with silicosis. The British journal of radiology. 2015;88(1049):20150028. doi: 10.1259/bjr.20150028. PubMed PMID: 25747897; PubMed Central PMCID: PMC4628485.

56. Faria ACD, Carvalho ARS, Guimarães ARM, Lopes AJ, Melo PL. Association of respiratory integer and fractional-order models with structural abnormalities in silicosis. Comput Methods Programs Biomed. 2019;172:53–63. Epub 2019/02/07. doi: 10.1016/j.cmpb.2019.02.003. PubMed PMID: 30902127.

57. Faria AC, Costa AA, Lopes AJ, Jansen JM, Melo PL. Forced oscillation technique in the detection of smoking-induced respiratory alterations: diagnostic accuracy and comparison with spirometry. Clinics. 2010;65(12):1295–304. PubMed PMID: 21340218; PubMed Central PMCID: PMC3020340.

58. Wang C, Ma J, Zhang S, Shao J, Wang Y, Zhou HY, et al. Development and validation of an abnormality-derived deep-learning diagnostic system for major respiratory diseases. NPJ Digit Med. 2022;5(1):124. Epub 2022/08/24. doi: 10.1038/s41746-022-00648-z. PubMed PMID: 35999467; PubMed Central PMCID: PMCPMC9395860.

59. Zhou Y, Mei S, Wang J, Xu Q, Zhang Z, Qin S, et al. Development and validation of a deep learning-based framework for automated lung CT segmentation and acute respiratory distress syndrome prediction: a multicenter cohort study. EClinicalMedicine. 2024;75:102772. Epub 2024/08/22. doi: 10.1016/j.eclinm.2024.102772. PubMed PMID: 39170939; PubMed Central PMCID: PMCPMC11338113.

60. Hyatt RE, Scandon, P.D., Nakamura, M. Interpretation of pulmonary function tests. Phyladelphia: Lippincott-Raven; 1997.

61. Ljung L. System identification: theory for the user: Prentice-Hall, Inc.; 1986.

62. Radics BL, Gyurkovits Z, Makan G, Gingl Z, Czövek D, Hantos Z. Respiratory Oscillometry in Newborn Infants: Conventional and Intra-Breath Approaches. Front Pediatr. 2022;10:867883. Epub 2022/04/22. doi: 10.3389/fped.2022.867883. PubMed PMID: 35444964; PubMed Central PMCID: PMCPMC9013809.

63. Lemes LN, Melo PL. Forced oscillation technique in the sleep apnoea/hypopnoea syndrome: identification of respiratory events and nasal continuous positive airway pressure titration. Physiol Meas. 2003;24(1):11–25. Epub 2003/03/15. doi: 10.1088/0967-3334/24/1/302. PubMed PMID: 12636184.

64. Sellares J, Acerbi I, Loureiro H, Dellaca RL, Ferrer M, Torres A, et al. Respiratory impedance during weaning from mechanical ventilation in a mixed population of critically ill patients. British journal of anaesthesia. 2009;103(6):828–32. doi: 10.1093/bja/aep301. PubMed PMID: 19887532.

65. Agustí A, Celli BR, Criner GJ, Halpin D, Anzueto A, Barnes P, et al. Global Initiative for Chronic Obstructive Lung Disease 2023 Report: GOLD Executive Summary. The European respiratory journal. 2023;61(4). Epub 2023/03/02. doi: 10.1183/13993003.00239-2023. PubMed PMID: 36858443.

66. András L, Dorottya C, Zoltán G, Gergely M, Bence R, Dóra B, et al. Airway dynamics in COPD patients by within-breath impedance tracking: effects of continuous positive airway pressure. European Respiratory Journal. 2017;49(2):1601270. doi: 10.1183/13993003.01270-2016.

